# Dual functions of *ZmGI1* in the photoperiodic flowering pathway and salt stress responses in maize

**DOI:** 10.1101/2021.05.12.443837

**Authors:** Fengkai Wu, Ling Liu, Yan Kang, Jing Li, Zhiyu Ma, Baba Salifu Yahaya, Jie Xu, Qingjun Wang, Xuanjun Feng, Jingwei Li, Erliang Hu, Yaxi Liu, Yanli Lu

**Author notes:** Correspondence and requests for materials should be addressed to Yanli Lu. These authors contributed equally to this work (Fengkai Wu, Ling Liu, Yan Kang).

## Abstract

The circadian clock perceives photoperiodic changes and initiates processes leading to floral transition. *GIGANTEA* (*GI*) primarily functions as a principal clock component that integrates environmental cues into regulation of growth and development in Arabidopsis. However, it is unclear whether *ZmGIs* regulate photoperiodic flowering and abiotic stress response. Here, we demonstrated that the expression of *ZmGI1* depicted a typical circadian pattern and was differentially expressed under LDs and SDs in photoperiodic sensitive and insensitive maize lines. The transcription level was significantly and positively correlated with days to silking and photoperiodic sensitivity in maize. Moreover, natural variation in *ZmGI1* was associated with maize photoperiod response and the fine-tuning of plant development traits. Overexpression of *ZmGI1*^Huangzao4^ induced early flowering and enhanced salt tolerance in Arabidopsis relative to the wild-type and *gi* mutants. ZmGI1 formed a protein complex with ZmFKF1 and acted as a positive regulator of flowering time by regulating *CONSTANS* transcription in the photoperiod pathway. The ZmGI1/ZmThox complex regulates oxidative stress induced by salt stress via a redox balance pathway. Over all, we have provided compelling evidence to suggest that *ZmGI1* is a pleotropic gene whose expression depicts a typical circadian rhythmic pattern and regulates flowering time and confers salt stress tolerance.

## Introduction

Plants precisely anticipate the onset of flowering by constantly monitoring environmental signals and coordinating endogenous cues to ensure a successful transition from the vegetative to reproductive growth stages. A complex network comprising various genetic and epigenetic regulators that responds to external stimuli and triggers floral transition has been well examined in the long day (LD) model species *Arabidopsis thaliana* (Blümel et al 2015). Daylength sensing (photoperiodism) is one of the most reliable seasonal cues that is exclusively measured by the circadian clock in vascular tissue (Shim et al 2017, Song et al 2015). The molecular mechanisms in the photoperiodic flowering pathway mainly involve three components: light input, circadian clock, and rhythm output (Shim et al 2017). Plants are known to perceive light signals (Day length, light quality, quantity, and direction) in mature leaves using various photoreceptors and transmit signals to the shoot apex to initiate flowering. Circadian clocks act as external time keeping mechanisms that modulate photoperiodism in plants (Creux & Harmer 2019, Millar 2004). Most components of the circadian clock are transcription repressors (Shim et al 2017) and there are multiple interconnected negative feedback loops that form a 24-h oscillator rhythm (Creux & Harmer 2019, Endo 2016, Inoue et al 2018, Locke et al 2006). The input signals from the ambient environment help reset the clock of the circadian system (Creux & Harmer 2019, Tóth et al 2001). Levels of the repressor and activator transcription factors in the circadian clocks are in constant flux, each peaking at a specific time of day and feeding back to regulate the expression of others (Creux & Harmer 2019, Shim et al 2017).

Circadian clocks integrate various environmental signals with endogenous cues to coordinate diverse physiological responses (Adams et al 2018, Inoue et al 2018, Qian et al 2014). In addition to its basic role in light and temperature modulation networks, the circadian clock also functions in multiple abiotic and biotic stress responses. GI is a unique plant specific nuclear protein involved in the circadian clock-regulated flowering pathway (Fowler et al 1999, Huq et al 2000, Mizoguchi et al 2005). *GI* plays a crucial role in regulating rhythm output and further increasing *CO* mRNA abundance; it also supervises the activity and stability of the CO protein, which regulates the accumulation of Flowering Locus T (*FT*) transcripts in phloem companion cells in leaves (Sawa & Kay 2011, Sawa et al 2007, Suárez-López et al 2001). The floral FT protein then moves from companion cells in the leaf phloem to the shoot apical meristem to promote flowering. In the photoperiodic flowering pathway, GI and FKF1 form a complex that degrades DOF factors, thus removing the inhibition of *CO* transcription, upregulating *FT* expression, and accelerating the time required to flower (Imaizumi et al 2005, Imaizumi et al 2003, Sawa et al 2007). There are two other mechanisms by which *GI* adjusts *FT* expression independent of *CO*. First, *GI* inhibits the expression of *TARGET OF EAT 1* (*TOE1*), thus upregulating *FT* transcription based on miR172 regulation (Jung et al 2007). GI also degrades the *FT* transcriptional repressors SHORTVEGETATIVEPHASE (SVP), TEMPRANILLO 1 (TEM1), and TEMPRANILLO 2 (TEM2), leading to high *FT* transcription (Sawa & Kay 2011). *GI* is therefore a major mediator between the circadian clock and the master regulators (*CO* and *FT*) in the photoperiodic flowering pathway.

Some studies have also implicated *GI* in other plant development functions, such as light signal perception (Oliverio et al 2007), cotyledon movement (Tseng et al 2004), and cell wall deposition (Edwards et al 2010). Additionally, *GI* functions as one of the crucial mediators that coordinates plant responses to various environmental stresses, such as cold (Fornara et al 2015, Fowler & Thomashow 2002), salinity (Kim et al 2016, Kim et al 2013), and drought (Riboni et al 2013, Riboni et al 2016). These results reveal that *GI* is involved in diverse biological processes of plant development and resistance to numerous stresses that threaten crop production. Information regarding the biological functions and regulatory mechanism of *ZmGIs* in the photoperiodic-dependent flowering pathway remains limited. Particularly, the function of *ZmGI*s in linking the photoperiodic pathway and stress resistance in maize is not well understood. In this study, we examined *ZmGI1* expression under both LDs and short days (SDs) in different maize lines. We collected genetic, biochemical, and physiological data to investigate the role of *ZmGI1* in regulating the photoperiodic flowering pathway and salt stress responses in maize. Based on the results of these analyses, we conducted further analysis on the function of ZmGI1 in promoting flowering under LDs.

## Materials and methods

### Plant materials, growth conditions, and flowering time investigation

Maize inbred line B73 was used to detect the expression patterns of *ZmGI1* in different tissues under field conditions in the spring of 2019 in Wenjiang, Sichuan, China. Six extreme phenotypic maize lines were selected from two association panels comprising 87 (Liu et al 2018) and 368 diverse core maize inbred lines (Li et al 2013), from China, USA, and CIMMYT included three tropical photoperiod-sensitive inbred lines (CMT-L189, CML202, and CML496) and three temperate-neutral/photoperiod-insensitive inbred lines (Mo113, CIMBL60, and Huangzao4). For the photoperiodic assays, the plants were transferred to a controlled growth chamber at 28 °C/light and 22 °C/dark for LDs (16 h light/8 h dark) or SDs (8 h light/16 h dark) and entrained for 2 weeks.

*ZmGI1* transgenic plants were generated by Pro_*35S*_::*ZmGI1*^Huangzao4^*-GFP* overexpressing in *Col*-0 through *Agrobacterium tumefaciens*-mediated transformation. T-DNA insertion *gi* mutant (CS879752) was obtained from the Arabidopsis Biological Resource Center. Plants were grown in greenhouse at 22–23 °C, 60% relative humidity, and under LD (16 h light/8 h dark) or SD (8 h light/16 h dark) conditions. Flowering time was measured by counting the number of days to bolting and/or the total number of rosette leaves when floral buds were visible (nearly 1 cm long) at the center of the rosette. A total of 20–24 plants were measured and averaged for each sample. Statistical significance was determined using a Student’s *t*-test (\*\**P* < 0.01; \**P* < 0.05).

### Association analysis

The polymorphisms among the 368 maize inbred lines were gathered from previous re-sequencing data (Li et al 2013) based on the physical location of *ZmGI1* in the maize genome. All lines were grown in three environments: one with LD (>13 h, Sichuan, SC) and two with SD (<12 h, Yunnan, YN and Guangxi, GX) growth conditions. Flowering-related traits were investigated and measured as days to anthesis (DTA), days to silking (DTS), days to tasseled (DTT), and anthesis-silking interval (ASI). The other characteristics of plant architecture, such as plant height (PH), ear height (EH), ear leaf length (ELL), and ear leaf width (ELW) and kernel traits including hundred kernel weight (HKW), kernel length (KL), kernel width (KW), and kernel thickness (KT), were described previously (Li et al 2013). The traits from plant architecture and kernel for each inbred line were calculated based on the best linear unbiased prediction (BLUP) to estimate the phenotype values, which were then used to implement an association analysis. The significant SNP variations in *ZmGI1* for the tested traits were calculated using TASSEL v5.0 software (Bradbury et al 2007) under a general linear model with a Q matrix indicative of population structure (GLM+Q).

### Subcellular localization

The ClonExpress^®^II system (Vazyme, Nanjing, China) was used to generate C-terminal enhanced green fluorescent protein (eGFP) fusion vector Pro_*35S*_::*ZmGI1*^Huangzao4^*-GFP*. Transient transformations of tobacco (*Nicotiana benthamiana*) leaves were performed using an agroinfiltration protocol that has been described previously (Wu et al 2016). In addition, we also transiently expressed the plasmid by PEG mediated in maize protoplasts. The fluorescence signals were detected, and images were acquired after 40 h of incubation at room temperature using a confocal microscope (LSM800; ZEISS, Oberkochen, Germany) with appropriate filters.

### Salt stress in Arabidopsis and maize

After germination on 1/2 MS medium 4 d, WT, *gi* mutant, and *ZmGI1* transformants were transplanted to 1/2 MS medium and with 100 mM NaCl, 250 mM mannitol, and 1/2 MS without phosphorus, various treatment conditions, respectively. After 12 d of cultivation, the photosynthetic capacity (ratio of variable to maximum chlorophyll fluorescence, *Fv/Fm*) was measured using a FluorCam 800MF system (Photon System Instruments, Drasov, Czech Republic) following the manufacturer’s protocol. When seedlings of six selected maize lines reached the three-leaf stage, plants were assigned to either a new nutrient solution under normal conditions or a nutrient solution with 250 mM NaCl for 7 d. Changes in the root morphology indices of treated seedlings were measured by a WinRhizo Pro 2008a image analysis system (Regent Instruments Inc., Quebec, Canada) equipped with a professional scanner (Epson XL 1000, Nagano, Japan). H_2_O_2_ accumulation in maize leaves and roots was quantified using a hydrogen peroxide assay kit (Beyotime, Shanghai, China) according to the manufacturer’s instructions. Statistical significance was determined using a Student’s *t*-test (\*\**P* < 0.01; \**P* < 0.05).

### Measurement of promoter activity of ZmGI1

To investigate the response of *ZmGI1* expression under salt stress, we fused the promoter of *ZmGI1* to a GUS reporter gene, and the recombinant transgenes were introduced into Arabidopsis to produce Pro_*ZmGI1*_::GUS transgenic plants. Leaves were infiltrated in GUS staining solution (50 mM sodium phosphate, pH 7.0, 10 mM EDTA, 0.1% Triton-100, 0.5 mg/ml X-gluc, 0.1 mM potassium ferricyanide, and 10% methanol) and incubated at 37 °C in the dark for 10–12 h, then washed in 70% ethanol several times until they were colorless to de-stain them before photographing. In addition, the activity of the *ZmGI1* promoter was evaluated and quantified by measuring the accumulation of *GUS*. Fresh leaves were collected at ZT9 from 12 d old seedlings that had been incubated for 24 h under either SD, LD, or LD plus 100 mM NaCl conditions. The expression levels of *GUS* were measured by RT-qPCR analysis using *IPP2* as relative control. Anti-GUS (Sigma, MO, USA) was employed to determine the accumulation of GUS protein using the western blot assay.

### Chlorophyll content estimation

To estimate changes in chlorophyll content in leaves under salt stress treatment in the selected six maize inbred lines, SPAD values measured using a portable chlorophyll meter (SPAD-502, Hangzhou Mindfull Technology Co., Ltd, China) represented relative chlorophyll contents. After 3 d of salt stress treatment, SPAD values for each unfolding leaf were measured 10 times at different leaf positions, and then mean values were calculated as an indicator of chlorophyll content. At least five plants from each line were measured, and statistical analyses were conducted using the data obtained from three independent experiments. Statistical significance was assessed via a Student’s t test (*P* ≤ 0.05).

### Proline content measurement

Free proline content was determined using a ninhydrin assay (Bates et al 1973). A total of 0.2 g fresh tissue from the six maize inbred lines was ground in liquid nitrogen, then 2 mL of 3% (w/v) sulfosalicylic acid was added to each sample at room temperature with constant shaking for 10 min to extract proline. Subsequently, the supernatant was obtained after centrifugation at 12,000 × *g* for 10 min. The supernatant was mixed with 2.5% (w/v) ninhydrin dissolved in glacial acetic acid and phosphoric acid, followed by boiling at 100 °C for 1 h. After rapid cooling and toluene extraction, the absorbance of the reaction mixture was measured at 520 nm with a microporous plate spectrophotometer (MQX200R2+Take3™, BioTek) according to the user’s manual. The proline content was calculated from the standard curve obtained using the proline standard (L-proline) solution and expressed based on fresh weight as µg g^-1^, with each experiment performed in triplicate.

### RNA extraction and RT-qPCR analysis

Total RNA was isolated from different tissues using a Plant Total RNA Isolation Kit (Foregene, Chengdu, China) according to the manufacturer’s instructions. cDNA synthesis was performed using 1 µg total RNA with a *TransScript*^®^II One-Step gDNA Removal and cDNA Synthesis SuperMix (TransGen, Beijing, China) with residual genomic DNA removed. The cDNA was diluted 5-fold with nuclease-free water and used as template for qRT-PCR analysis, which was performed with three technical replicates using a *TransScript*^®^II Green One-Step qRT-PCR SuperMix (TransGen, Beijing, China) and the expression of housekeeping genes *ZmUBI1* in maize and *IPP2* in Arabidopsis were used as internal controls. The primers used are listed in Table S1.

### Yeast two-hybrid cDNA library screening and confirmation

The CloneMiner^™^ II cDNA library construction kit (Invitrogen) was employed to construct a cDNA library in P178 maize seedlings. High-quality cDNA libraries were constructed into pGADT7 (AD) vector and transformed into Y187 competent yeast cells by OEbiotech (Shanghai, China). Yeast two-hybrid (Y2H) library screening was performed using the Clontech two-hybrid system according to the manufacturer’s instructions. The constructed carrier, Y2HGold competent yeast cells with pGBKT7-ZmGI1 (BD-ZmGI1), was applied to screen the P178 cDNA library after it was tested for auto-activation as a bait vector. The transformants were screened on SD/-Ade/-Leu/-Trp/-His/X-α-Gal (Coolabar, Beijing, China) agar plates and incubated for 2–4 d at 28 °C. Prey plasmids were extracted and sequenced from single blue colonies, which are putatively positive clones.

To further confirm interactions, candidate genes from positive clones were inserted into BD vectors and their interaction abilities were verified by co-transformation with AD-ZmGI1 into Y2HGold strains. pGBKT7-53 and pGBKT7-Lam were co-transformed with pGADT7-T as positive and negative controls, respectively. Transformants were plated and cultured on SD/-Trp/-Leu and SD/-Ade/-Leu/-Trp/-His/X-α-Gal agar plates to test for interactions.

### Split luciferase (LUC) complementation

The full-length coding sequences of *ZmGI1* and *ZmFKF1a* were amplified by the specific primers listed in Table S1. The PCR products were cloned into nLUC/ cLUC vectors via the ClonExpress^®^II system (Vazyme, Nanjing, China). GV3101 harboring the corresponding constructs were resuspended in an injection infiltration buffer (10 mM MgCl_2_, 10 mM MES pH 5.6, and 100 µM acetosyringone). For co-infiltration, equal volumes of two different strains carrying the indicated nLUC and cLUC constructs were mixed and infiltrated into *N. benthamiana*. After 48 h, the infiltrated leaves were sprayed with luciferin, and fluorescence was detected in the dark using a CCD camera.

### In vivo co-immunoprecipitation

To further verify the interactions between ZmGI1 and ZmFKF1a *in vivo*, we co-infiltrated the Agrobacterium strains carrying the ZmFKF1a-Flag or ZmTHOX-mCherry and ZmGI1-GFP plasmids into 4-week-old *N. benthamiana* leaves. For co-immunoprecipitation (Co-IP) assays, all plant tissues were ground in liquid nitrogen, proteins were extracted, and IP procedures were performed at 4 °C. Proteins were extracted in lysis buffer [50 mM Tris-HCl pH 7.5, 150 mM NaCl, 5 mM EDTA, 2 mM DTT, 0.1% Triton X-100, 1 mM phenylmethyl sulfonyl fluoride (PMSF), and complete protease inhibitor cocktail tablets (Roche, Basal, Switzerland)] for 30 min. The remaining supernatant was incubated with Protein G-coupled magnetic beads (Sigma, MO, USA) that captured with anti-GFP for 4 h. The beads were then washed three times with 500 µL of lysis buffer with protease inhibitors after adsorbing on the magnetic frame for 1 min. The bead-precipitated proteins were eluted with 2× SDS loading buffer at 95 °C for 10 min. The ZmGI1-GFP, ZmFKF1a-FLAG, and ZmTHOX-mCherry proteins were detected by western blot using anti-GFP (Sangon Biotech, Shanghai, China), anti-Flag (Sigma, MO, USA), and anti-mCherry (Proteintech, Wuhan, China) antibodies, respectively.

## Results

### Natural variations in *ZmGI1* significantly associated with maize photoperiod sensitivity

The expression of two identified *GIGANTEA* (*ZmGI1*: Zm00001d008826, *ZmGI2*: Zm00001d039589), based on CuffLinc FPKM values of inbred line B73 (12 h day/12 h night) (Lai et al 2020), showed that *ZmGI1* exhibited much higher than *ZmGI2* in maize (Figure S1A), suggesting that *ZmGI1* plays a primary role in regulating the photoperiodic flowering pathway. In addition, *ZmGI1* is extensively expressed in different tissues, especially in roots, stems, and leaves (Figure S1B). The results suggested that *ZmGI1* might be involved in multiple biological processes.

The significant variation in phenotypes of flowering-related traits in maize is due to the latitudes of four planting environments vary considerably. Meanwhile, the agronomic traits among maize association panel were observed. Based on the physical location of *ZmGI1* in the maize genome, we identified 23 polymorphic sites in coding sequence with a minor allele frequency (MAF) ≥ 0.05. The association between natural variation in *ZmGI1* and 12 traits (DTA, DTS, DTT, ASI, HKW, KL, KW, KT, PH, EH, ELL, and ELW) in four different environments (Table S2) was investigated, and three SNP (S7632, S7960, and S8002) located in coding regions were simultaneously significantly associated with DTA and DTS under multiple environments at *P* < 0.01 (Figure 1A, B; Table S2), indicating that the sites have crucial roles in plant flowering. In addition, eight SNP were simultaneously significantly associated with HKW and KL (Figure 1C; Table S2). Three sites located in a complete LD block were significantly associated with PH and ELW, explaining 3.6% and 3.8% of phenotypic variation, respectively (Figure 1D; Table S2). Detailed information on the location, genotype, frequency, and statistical value of each site is presented in Table S2. The results indicate that *ZmGI1* is associated with flowering time in maize development.

**Figure 1.**
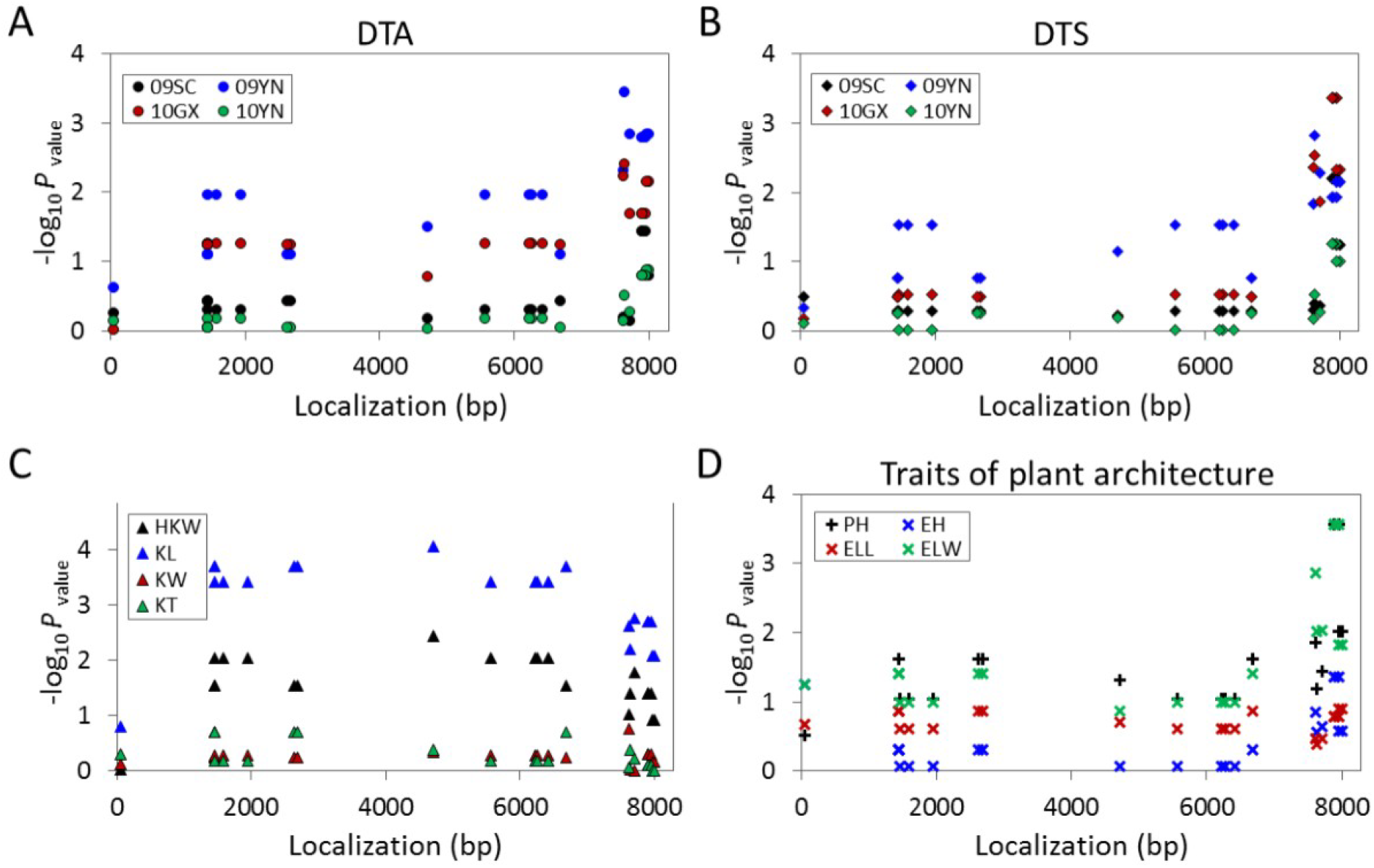
Natural variations within *ZmGI1* are associated with agronomic traits. **(A, B)** Natural variation in *ZmGI1* associated with day to anthesis (DTA) and day to silk (DTS) under various photoperiod conditions, respectively. Each circle and rhombus represent a polymorphic SNP. 09SC, 09YN, 10GX, 10YN indicate 2009 in Sichuan, 2009 in Yunnan, 2010 in Guangxi and 2010 in Yunnan, respectively. **(C, D)** Natural variation in *ZmGI1* associated with traits of kernel and plant architecture, respectively. All the tested traits were performed under four photoperiod conditions and the BLUP values of traits were calculated to assess the association between *ZmGI1* and agronomic traits, like hundred kernel weight (HKW), kernel length (KL), kernel width (KW), kernel thickness (KT), plant height (PH), ear height (EH), ear leaf length (ELL), and ear leaf width (ELW). Each triangle and cross represent a polymorphic SNP.

### Circadian clock regulates the expression of ZmGI1 in maize

The two *ZmGIs* exhibited a 24 h rhythmic expression that peaked at ZT9 or ZT12 (Figure S1A), indicating maximum accumulation of *ZmGI* mRNAs in the early evening (Lai et al 2020). Obvious differences in *ZmGI1* expression among maize inbred lines under the LD or SD conditions were revealed (Figure 2A, B). Generally, the photoperiod-sensitive tropical inbred lines exhibited significantly higher levels of *ZmGI1* expression than those of the temperate lines, especially under the LD condition. Expression peaks appeared at ZT9 in tropical lines and ZT12 in temperate lines under LD conditions, indicating a distinction in regulation between the two germplasm groups. Therefore, the tropical and temperate germplasms could be well distinguished based on the expression patterns under LDs, suggesting a close relationship between the expression of *ZmGI1* and photoperiodic sensitivity in maize. To further confirm this relationship, we calculated the correlation coefficient of peak expression values and photoperiodic flowering-related traits. The *ZmGI1* mRNA accumulation among the lines was significantly positively correlated with DTS and photoperiodic sensitivity under both LD (r = 0.91, *P* < 0.05) (Figure 1C) and SD conditions (r = 0.85, *P* < 0.05) (Figure 1D), implying that *ZmGI1* might play a vital role in photoperiodic sensitivity regulation. Furthermore, the results from semi-quantitative PCR performed in the six maize inbred lines under LD conditions indicated a strict diurnal cycle and expression regularity, which is consistent with these findings (Figure S1C).

**Figure 2.**
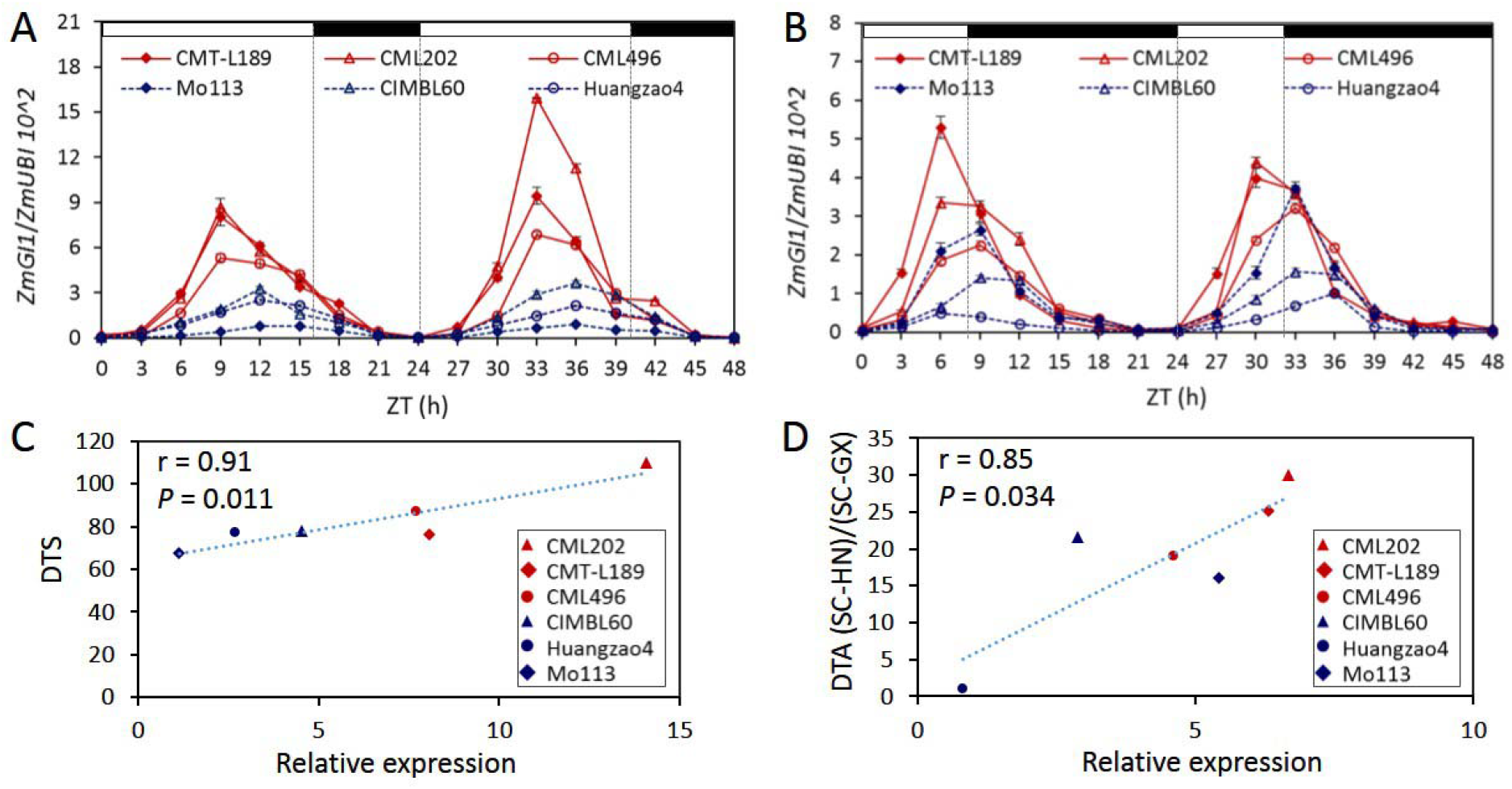
Expression characteristics of *ZmGI1* in maize. Diurnal expression of *ZmGI1* in maize under LDs **(A)** and SDs **(B)** conditions. Samples were harvested every three hours during a 48-h period. Red and blue lines indicate photoperiodic sensitive and insensitive maize lines, respectively. **(C)** The correlation between the expression of *ZmGI1* in six maize inbred lines and the Day to Silk (DTS) strait under long-day condition in Sichuan at 2009 (Day length >13.5 h). **(D)** The correlation between the expression of *ZmGI1* and the photoperiod sensitivity index calculated from Day to Anthesis (DTA) in the long-day condition of Sichuan at 2009 (Day length >13.5 h) and the short-day of Hainan at 2009 (Day length <12 h) and Guangxi at 2010 (Day length <11.5 h).

ZmGI1-GFP was simultaneously co-localized in the nucleus (RFP marker) and cytoplasm in *N. benthamiana* leaves (Figure S1D), even though GI is reported as a nuclei protein in Arabidopsis (Fowler et al 1999). ZmGI1 protein was detected in the nucleus and cytoplasm of maize protoplast cells, with GFP (Pro_*35S*_::*GFP*) used as a control (Figure S1E), indicating that the ZmGI1 was localized in the nucleus and cytoplasm.

### Overexpressing ZmGI1 promotes flowering under LD conditions in Arabidopsis

Three independent homozygous transgenic lines (OE#8, OE#10, and OE#14) showing increased accumulation of *ZmGI1* mRNA from a T_4_ population, confirmed to contain a single Mendelian locus, were used to investigate the character of responsiveness to day length (Figure 3A, B). Under LDs, the flowering time of transgenic lines was approximately 7 d earlier than that of the WT (Figure 3B, C). Under SDs, flowering time was 3 d earlier in the transgenic plants compared to that in the WT. In comparison, flowering in the *gi* mutants was delayed by approximately 21 d compared to that in the WT under both photoperiodic conditions. Meanwhile, the number of rosette leaves in transgenic plants was almost indistinguishable from that of the WT under LD conditions (Figure 3D, E). However, the number of rosette leaves in the *gi* mutants was significantly higher than that of WT and transgenic lines. These results indicated that delayed flowering under both photoperiodic conditions in the *gi* mutants was due to a prolonged floral transition, whereas overexpression of *ZmGI1* contributed to hastened floral transition.

**Figure 3.**
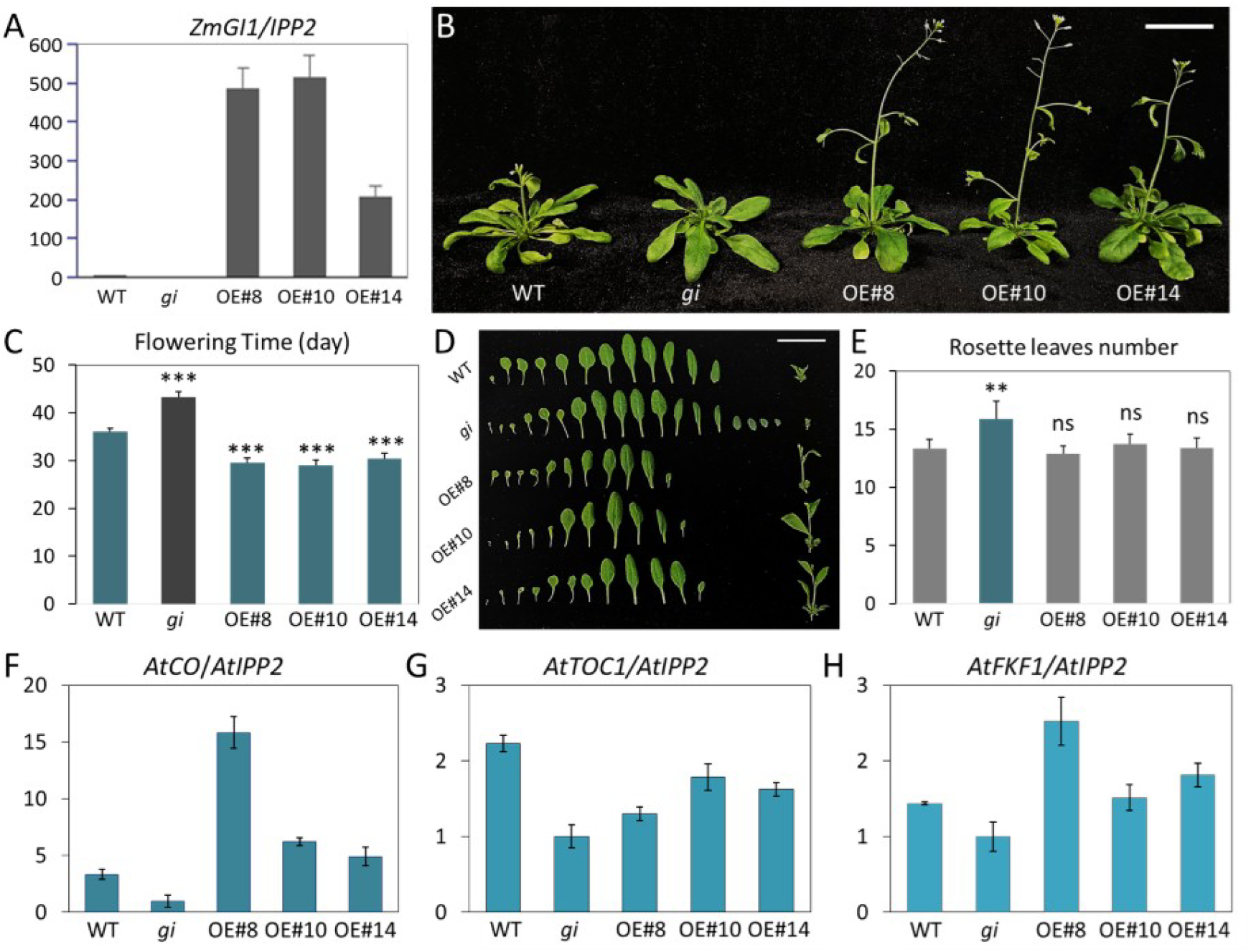
Phenotypes of overexpression *ZmGI1* in Arabidopsis under LDs. **(A)** The transcription of *ZmGI1* was detected in 36-d-old seedlings of different overexpression lines at ZT10. (B) Phenotypes of different genotypic plants after 36 d treatment under LDs grown in soil. **(C)** Bolting time assay in different genotypes under LDs. Values are means ± SD (24 ≥ n ≥20); \*\**P* < 0.01 (Student’s t test). **(D)** Morphological phenotype of the leaves of 30-d-old WT, *gi* and *ZmGI1* overexpression plants grown in soil under LDs. **(E)** Number of rosette leaves calculated from d; **(F-H)** Expression Patterns of *AtCO* (f), *AtTOC1* (g), and *AtFKF1*(h) in wild-type, *gi*, and overexpression plants under LDs. The expression levels measured by RT-qPCR are shown relative to the housekeeping gene *IPP2*. Each plant sample rosette leaves were harvested in 35 d after germination at ZT10, a relatively high expression point of *ZmGI1*. ***, *P* < 0.001; ns, not significant (Student’s *t* test). The scales are 3 cm in (A) and (D).

Nine genes involved in the photoperiodic flowering pathway, excluding *AtCDF1* and *AtFT*, were detected at the selected sampling points. All the tested genes exhibited relatively low transcript levels in the *gi* mutants (Figure 3 and S2). The level expression of florigene *AtCO* increased significantly in the overexpressed plants under LD conditions (Figure 3F), indicating that *ZmGI1* had a considerable effect on *AtCO* expression and functions upstream of *AtCO* in the LD photoperiodic flowering pathway. However, *AtTOC1* and *AtPRR7* were downregulated in overexpression plants (Figure 3G and S2), suggesting that *ZmGI1* was involved in the feedback loop in the evening. This result suggests that *ZmGI1* upregulates the expression of *AtCO* and promotes flowering in the photoperiodic pathway in Arabidopsis. The difference in *ZmGI1* expression in the regulation of *AtFKF1* and *AtTOC1* suggested that *ZmGI1* had a specific model of supervising flowering time owing to the heterogeneous overexpression or the dissimilar function of *ZmGI1* in maize.

### ZmGI1 responds to photoperiod and salt stress

*GI* is primarily involved in circadian rhythms and flowering time regulation and also regulates diverse physiological processes in Arabidopsis. To gain insight into the role that *ZmGI1* function in regulating abiotic stress response, we examined the root morphological traits of the WT, *gi* mutants, and *ZmGI1* transgenic lines under phosphorus starvation, drought stress, and salt stress. All the tested genotypes showed normal root elongation and equivalent lateral roots on 1/2 MS medium and manifested similar degrees of growth inhibition under phosphorus deficiency and drought stress (Figure S3A). Interestingly, *gi* mutants were highly hypersensitive to salt stress, manifesting much slower root elongation and fewer lateral roots on 1/2 MS medium supplemented with NaCl compared to the WT (Figure S3A, B).

To investigate whether the activity of the *ZmGI1* promoter is regulated by photoperiod and salt stress, three transgenic lines in which the *ZmGI1* promoter was fused with a *GUS* reporter gene were generated and detected by GUS staining and expression analysis in Arabidopsis. The GUS staining results showed that there were significant differences in expression under SDs, LDs, and LDs with salt stress (Figure 4A). Compared with the colorless control of the wild type, the transgenic lines had a deeper color under SDs than LDs, which indicated that the promoter of *ZmGI1* had more powerful activity under SDs (Figure 4A-C). Moreover, salinity treatment observably enhanced the activity of the *ZmGI1* promoter under LDs, even more strongly than that under SDs (Figure 4A, B). The results of the transcriptional analysis also showed that the promoter had the highest activity under salt stress transfer from SDs to LDs (Figure 4B). The observed levels of accumulated GUS protein were consistent with their transcription levels under LDs, SDs, and LDs with salt stress (Figure 4B-D). Interestingly, GUS protein accumulation was higher under SDs than under the SD plus salinity treatment (Figure 4E). Salt stress enhanced the activity of the *ZmGI1* promoter and promoted GUS protein accumulation under LDs but reduced expression under SDs. Collectively, these results suggest that the activity of the *ZmGI1* promoter was regulated by both photoperiod and salt stress.

**Figure 4.**
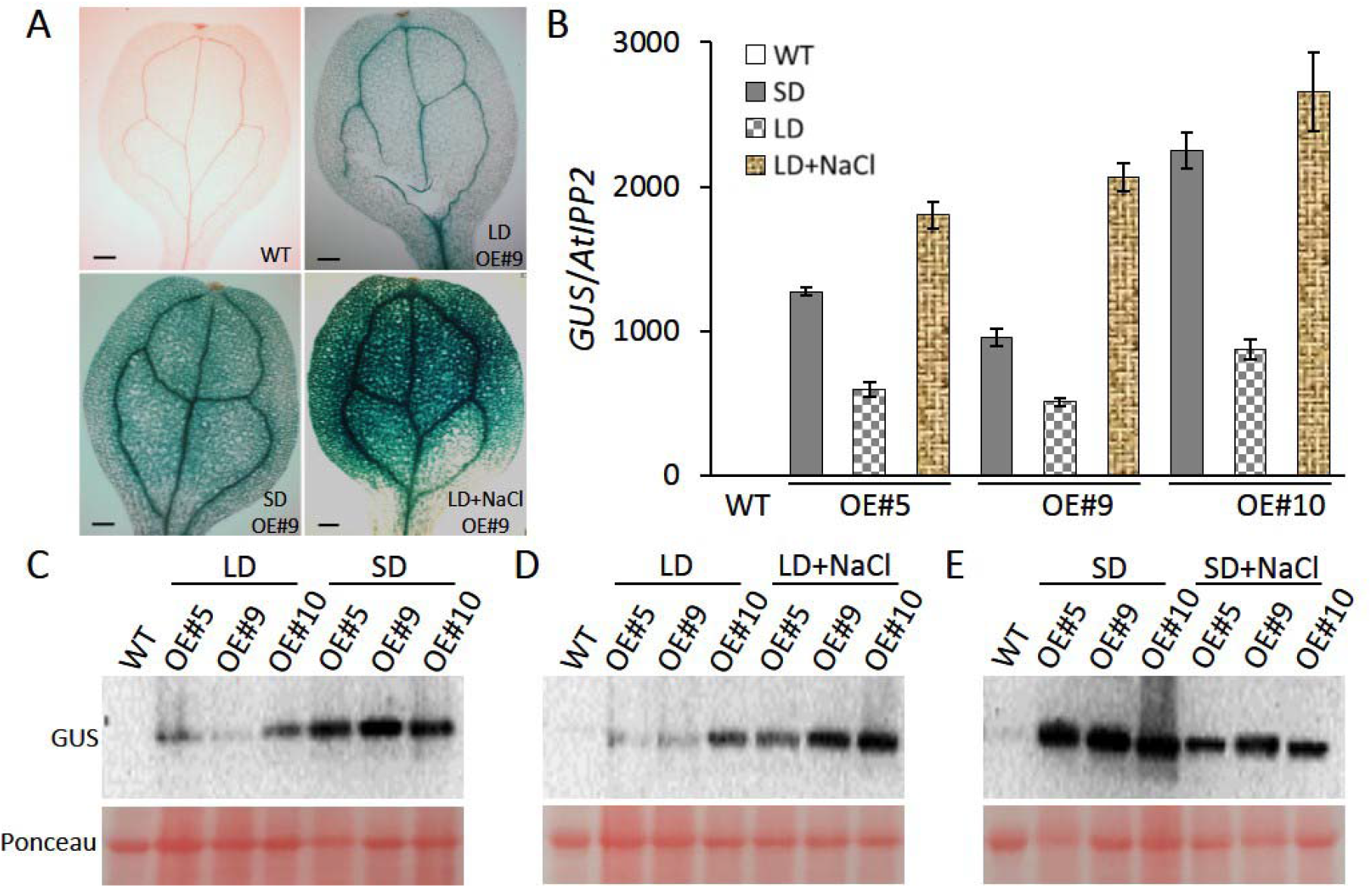
Expression analysis of transcription and protein level in Pro_*ZmGI1*_::GUS transgenic lines in Arabidopsis. **(A)** GUS expression of mature leaves were stained in transgenic seedlings. The 7-d-old transgenic lines were cultured on 1/2MS in SD, and then transferred to LD and with salt stress medium for 24 h. Scales are 200 μm. **(B)** The relative transcriptional levels of *GUS* were detected in different genotypes of Arabidopsis from (A). **(C-E)** Comparative determination of protein levels in LDs and SDs (C), LDs and salt stress (D), SDs and salt stress (E) used with anti-GUS in different genotypes of Arabidopsis. Similar results were obtained in three independent biological repetitions.

### Overexpression of *ZmGI1* enhances salt tolerance in Arabidopsis

Intuitively, overexpression lines exhibited significantly higher salt stress tolerance compared to WT and mutant genotypes (Figure 5A-B), with marked superiority in terms of root morphological traits such as total root length, number of lateral roots, root volume, and root surface area (Figure 5D-G). These results indicate that *ZmGI1* plays a crucial role in root elongation and lateral root formation under salt stress. Meanwhile, an increase in the transcriptional level of *ZmGI1* was observed under NaCl treatment in the overexpression lines over time; e.g., the transcription level of *ZmGI1* was 14 times higher than that of 0 h after salt treatment for 9 h (Figure 5C). This result indicates that salt stress induces the accumulation of *ZmGI1* mRNA or inhibits its degradation, and *ZmGI1* improves the tolerance of Arabidopsis to salt stress through post-transcriptional regulation.

**Figure 5.**
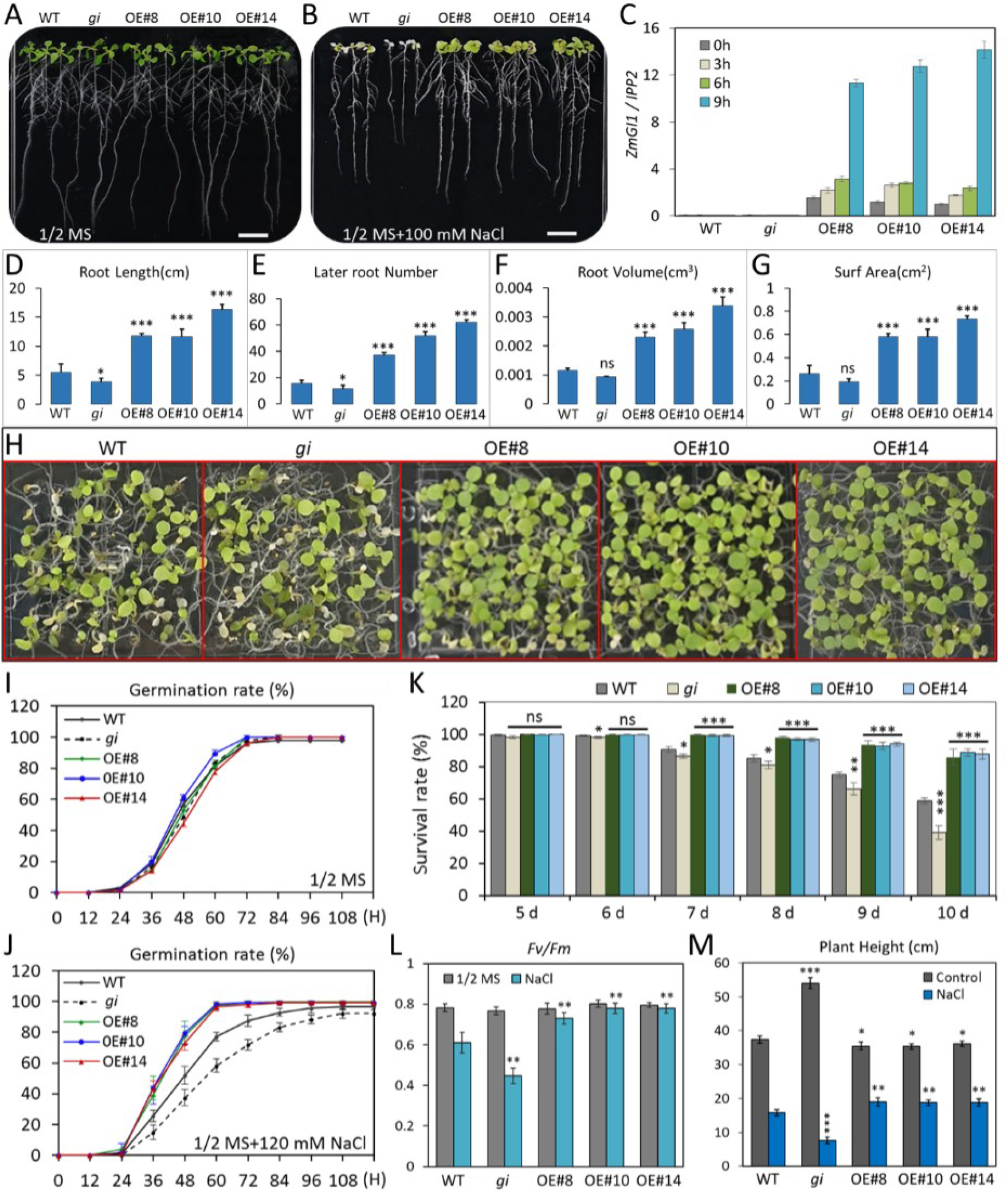
Overexpression of *ZmGI1* enhances salt tolerance in Arabidopsis seedlings. **(A, B)** Phenotype of WT, *gi* mutant and transgenic plants of *ZmGI1* germinated on 1/2 MS and 1/2 MS with NaCl (100 mM) medium. Four-day-old seedlings of different genotypes grown on 1/2 MS medium were transferred to new 1/2 MS and 1/2 MS with NaCl (100 mM) medium. Photograph was taken 5 d and 7 d after the transfer to 1/2 MS and 1/2 MS with NaCl (100 mM) respectively. Scale is 1 cm. **(C)** The expression levels of *ZmGI1* in different genotypes of Arabidopsis; **(D-G)** The root morphological traits of WT, *gi* and overexpressed *ZmGI1* lines under 1/2 MS with 100 mM NaCl stress. Values are means ± SD (n = 8). **(H)** Effect of salt stress shock on plant survival. WT, *gi* mutant and transgenic plants of *ZmGI1* germinated on 1/2 MS with NaCl (120 mM) media. Photograph was taken 10 d after treatment. **(I-J)** Germination rates of WT, *gi* mutant and over-expressed *ZmGI1* lines on 1/2 MS and under salt stress, respectively. Values are means ± SD (n = 5), 64 individual seedlings in each repeat. **(K)** Effect of salt stress on plant survival. WT, *gi* mutant and transgenic plants of *ZmGI1* were sown on 1/2 MS with NaCl (120 mM) medium. Survival rates of seedlings at the indicated time point after germination were calculated. Values are means ± SD (n = 5). **(L)** The effect of the maximum quantum efficiency (*Fv/Fm*) of PSII changes on salt stress in WT, *gi* mutant and transgenic plants of *ZmGI1*. Seedlings were grown for 12 d under continuous light with salinity treatment (120 mM NaCl) and then shifted to the dark for 2 h before PSII measurement. 64 individual seedlings of different genotypes in one culture dish were divided into a group and generated mean vales of *Fv/Fm*. The results represent the mean and SD of *Fv/Fm* measurements of three times repeats. **(M)** Effects of plant height at mature stage in WT, *gi* mutant and over-expressed *ZmGI1* lines under salt stress. The seeds were sown on 1/2 MS and transferred to soil and treatment with 150 mM NaCl at the full life cycle of Arabidopsis. Plant height were calculated from at least 8 individual seedlings. *, *P* < 0.05; **, *P* < 0.01; ***, *P* < 0.001, ns, not significant (Student’s *t* test).

The rates of seed germination and cotyledon greening in the *gi* mutants were normal on 1/2 MS medium, but, under 120 mM NaCl, their growth was retarded, and they experienced significantly higher mortality than WT (Figure 5H-K). Contrarily, under 120 mM NaCl, the transgenic lines exhibited superior growth, and *ZmGI1* overexpression promoted plant survival under salt stress (Figure 5 H-K). The maximum efficiency of PSII (*Fv/Fm*) was measured to assess the photosynthesis activities of *gi* mutants and transgenic plants. A decrease in the average value of *Fv/Fm* in WT and *gi* (Figure 5Land S3B) was detected, probably due to photodamage or downregulation of PSII reaction centers from salt stress. A consecutive salinity treatment throughout the life cycle of Arabidopsis resulted in yield reduction in the WT and higher lethality in the *gi* mutants under LDs compared to the overexpression lines (Figure 5K and S3D). A novel function of *GI*/*ZmGI1* in controlling plant height was found, and significant variation in plant height among the different tested genotypes was observed under both control and salt stress conditions (Figure 5M and S3C-D). Under normal conditions, *GI*/*ZmGI1* acted as an inhibitor of plant height, with the *gi* mutant becoming taller than transgenic plants, which exhibited dwarfish phenotypes. This outcome, which was opposite that under the salt stress treatment, suggested that *ZmGI1* has an advantageous physiological function in maintaining yield under saline stress. The plants on the NaCl and 1/2 MS medium containing high concentrations of mannitol, which can induce osmotic stress but not ion toxicity, had indistinguishable roots among all genotypes. The results confirmed that the salt hypersensitive phenotype of the *gi* mutant was caused by ion toxicity. Hence, overexpression of *ZmGI1* in Arabidopsis contributed remarkably to improved salt tolerance.

### Photoperiodic insensitivity improves salt tolerance in maize

To understand whether *ZmGI1* enhances salt tolerance in maize, three inbred lines from the photoperiod-sensitive maize panel and three inbred lines from the photoperiod-insensitive maize panel were subjected to a salinity treatment. Interestingly, we observed that the photoperiod-insensitive inbred lines exhibited significantly enhanced salt tolerance (Figure 6A). Proline is one of the protective compounds that help plants acclimatize to various stresses; to determine the relationship between salt tolerance and proline content in maize leaves and roots, we examined the proline levels under both normal conditions and salinity stress conditions. Generally, the proline content in photoperiod-sensitive inbred lines was significantly higher than that in the insensitive inbred lines (Figure S4A). In addition, the proline content significantly increased in the tissues of all inbred lines after salt stress treatment. Among these, the proline level in the photoperiod-insensitive inbred line of Huangzao4 was upregulated eight-fold (Figure 6B and S4A). Salt stress resulted in increased proline accumulation in the leaves and roots in the three photoperiod-insensitive maize lines with extreme salt tolerance, but it increased only in the leaves of the photoperiod-sensitive lines (Figure 6B and S4A).

**Figure 6.**
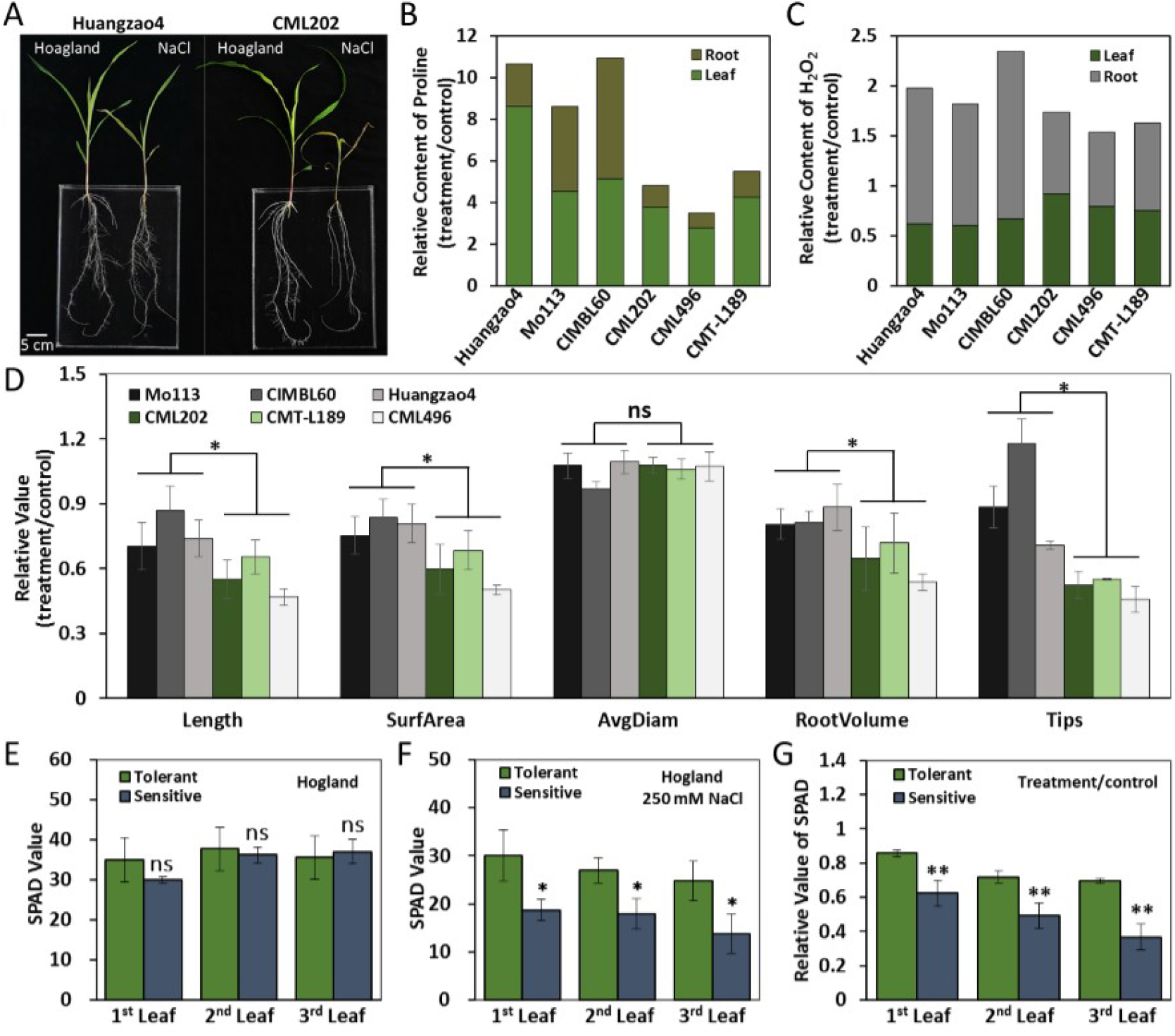
Phenotype of seedlings under salt stress in maize. **(A)** Effects of salinity on maize growth. The extremely salt tolerant and sensitivity maize lines were cultivated with Hoagland solution under neutral photoperiod (light 12 h/12 h dark) for 17 d and transferred to salt stress with 250 mM NaCl. Photographs were taken 7 d after transferred. Scales are 5 cm. **(B, C)** Relative content of proline (B) and H_2_O_2_ (C) in maize seedlings after 7d salt stress, respectively. The ratios were calculated from the treatment and control under both roots and leaves. **(D)** The relative variation ratio of root morphological traits in maize under 250 mM NaCl stress to control. n = 8 per column. **(E-G)** The chlorophyll content of salt tolerant and sensitive lines after 3 d salinity treatment.

Accumulations of H_2_O_2_ in leaves and roots under two treatments were quantified by biochemical testing. There was a slight difference in the accumulation of H_2_O_2_ between the photoperiod-sensitive and -insensitive lines, and there was also a much higher H_2_O_2_ content in the leaves than in the roots; however, salt stress decreased total H_2_O_2_ content (Figure S4B). Under the salt stress treatment, the H_2_O_2_ contents of leaves and roots in the three sensitive lines were slightly decreased (Figure 6C and S4B). Interestingly, there was a greater difference in the reduction of leaf H_2_O_2_ content in insensitive lines than in sensitive lines after salinity treatment (Figure 6C). In particular, it should be noted that the H_2_O_2_ content in the roots of insensitive lines increased under salt stress, the ratios of H_2_O_2_ content in roots between the salinity treatment and control treatment were much higher than in the sensitive lines (Figure 6C and S4B). The results indicate that although H_2_O_2_ was mainly concentrated in the leaves, the increase in root H_2_O_2_ content under salt stress treatment facilitated salt tolerance in the maize inbred lines.

In the three extremely salt-tolerant, photoperiod-insensitive inbred lines, the salinity treatment: control treatment response ratios for root length, surface area, root volume, and total root tips were notably increased compared to those of the photoperiod-sensitive lines (Figure 6D). We further compared the SPAD values of salt-tolerant and salt-sensitive plants under both control and stress conditions. There were no obvious differences among the three tested unfolding leaves under normal conditions (Figure 6E). However, a significant difference in the SPAD values was identified between the sensitive and insensitive genotypes under salt-stress treatment (Figure 6F), and this indicates that chlorophyll concentration could be a key factor for evaluating salt tolerance in maize. Moreover, the third leaf showed a huge reduction in chlorophyll concentration (Figure 6E-G), indicating that the chlorophyll in the old leaves degraded earlier than that in the younger ones.

### ZmGI1 interacts with ZmFKF1a and ZmTHOX

To elucidate the molecular mechanisms underlying the function of *ZmGI1* proteins in flowering time regulation and other biological functions, we performed a yeast two-hybrid assay using ZmGI1 as bait to identify the potential interacting proteins. The transcriptional activation activity of ZmGI1 in Y2H Glod was detected with full-length and truncated fragment constructs (amino acids 1 to 358, 359 to 612, 613 to 804, and 805 to 1163) (Figure S5). The truncated fragment of the ZmGI1 protein with no transcriptional activation activity (amino acids 359 to 1163) was used as bait to screen the yeast prey cDNA library prepared from whole P178 maize plants. The complete open reading frame of three candidate genes, detected in more than four independent clones, were revealed to interact with ZmGI1^359-1163^. None of the three proteins showed transcriptional activation activity (Figure 7A). Subsequently, Y2H assay demonstrated ZmFKF1a (Zm00001d007445), ZmSKIP35 (Zm00001d016858), and ZmTHOX (Zm00001d018461) interacted with ZmGI1 (Figure 7B). The interaction between FKF1 and GI is known to regulate flowering time in Arabidopsis (Imaizumi et al 2005), hence ZmFKF1a and ZmGI1 might perform similar functions in maize. *SKIP* genes are known for their function in alternative splicing of transcripts (Cui et al 2017, Wang et al 2012), suggesting that *ZmGI1*’s many alternative splicing activities could have resulted from its physical interaction with ZmSKIP35. Most biotic and abiotic stresses in plants are associated with redox reactions (Lázaro et al 2013). Thus, we assumed that the overexpression of *ZmGI1* regulates the Trx system by recruiting the ZmTHOX protein to improve stress tolerance.

**Figure 7.**
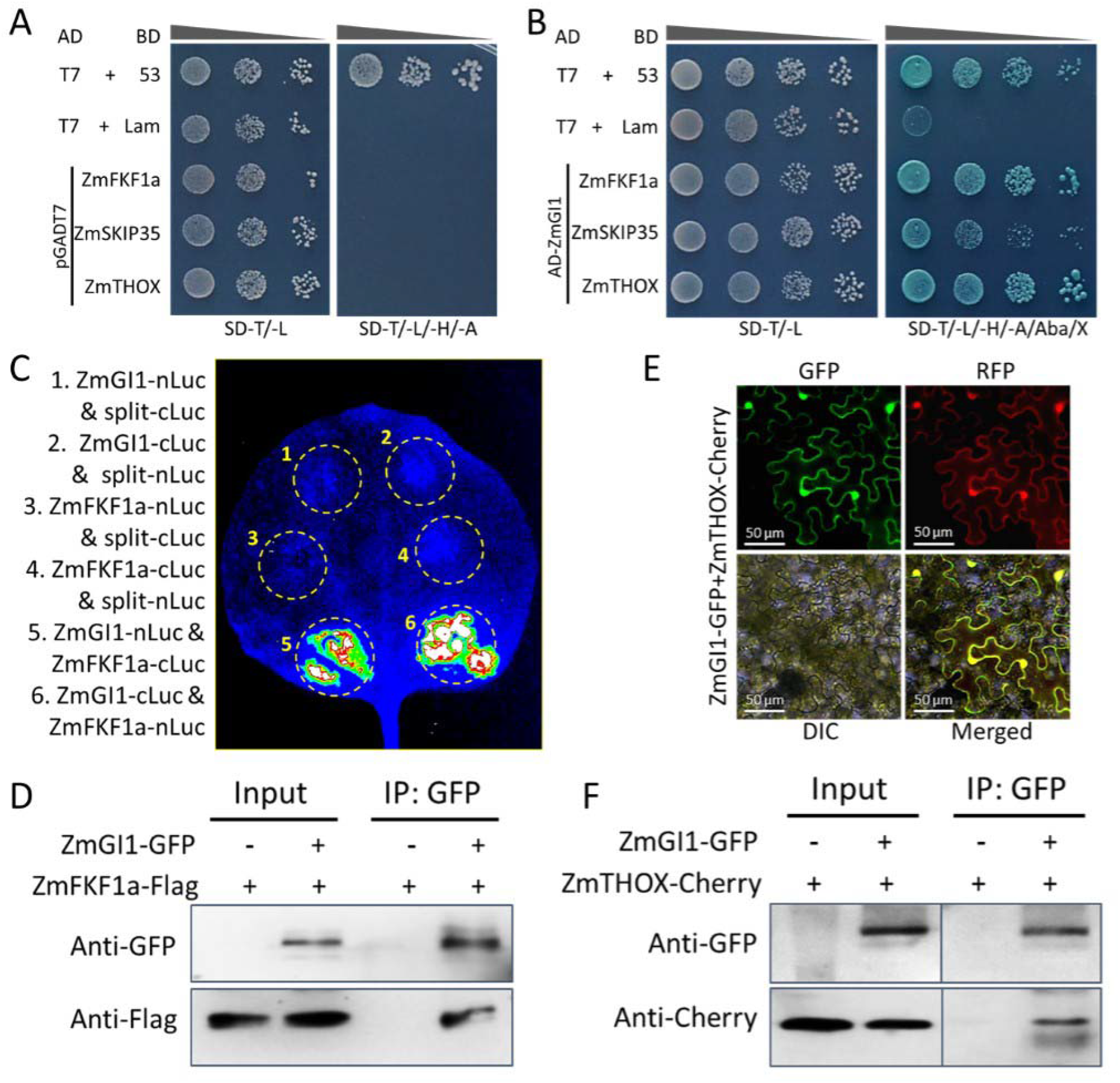
Interactions of ZmGI1 with ZmFKF1and ZmTHOX. **(A)** Self-activation verification of the candidate proteins. **(B)** ZmGI1 interacts with library screened proteins in yeast two-hybrid assay. **(C)** ZmGI1 interacts with ZmFKF1a in split-luciferase complementation (split-LUC) assay. The constructs to express the indicated fusion proteins were transformed to *N. benthamiana* leaves through Agrobacterium infiltration. Luciferase activity was determined at 48 h after infiltration. Three independent repeats were with consistent results. **(D)** ZmGI1 interacts with ZmFKF1a in tobacco by Co-IP. Coimmunoprecipitation assay (Co-IP) showing the interaction between ZmGI1 and ZmFKF1a. ZmGI1-GFP and ZmFKF1a-Flag were expressed in *N. benthamiana*. Immunoprecipitation was performed by using antibody anti-GFP. Immunoblottings were conducted using anti-GFP and anti-Flag antibodies. **(E)** Co-localization of the ZmGI1 and ZmTHOX in *N. benthamiana* leaves. ZmGI1 and ZmTHOX were fused with GFP and mCherry, respectively. Bars = 50 µm. **(F)** ZmGI1 interacts with ZmTHOX in tobacco by Co-IP assay. Plasmids containing ZmGI1-GFP and ZmTHOX-mCherry were co-transformed into tobacco leaves. Anti-GFP magnetic beads were used to immunoprecipitate the proteins, which were further analyzed by immunoblotting with anti-GFP and anti-mCherry antibodies.

Split luciferase (split-LUC) complementation assay was used to confirm whether ZmFKF1a interacts with ZmGI1 in the abaxial epidermal cells of tobacco leaves. Our results demonstrated that ZmFKF1a and ZmGI1 physically interact *in vivo* (Figure 7C). Coimmunoprecipitation (Co-IP) assays using *N. benthamiana* leaves further confirmed the interactions between ZmFKF1 and ZmGI1 (Figure 7D), suggesting that ZmFKF1a and ZmGI1 do exist in a complex. A further Co-IP assay by co-expressing ZmGI1 and ZmTHOX proteins in *N. benthamiana* leaves also confirmed the interaction between these two proteins (Figure 7F). In addition, ZmGI1 and ZmTHOX were fused with green fluorescence (ZmGI1-eGFP) and mCherry (ZmTHOX-mCherry), respectively. The co-expression results showed that both proteins were localized in the nuclei and cytoplasm in *N. benthamiana* leaves (Figure 7E). The consistent localization of these two proteins makes their physical interaction more credible.

## Discussion

### The circadian clock was visibly different in photoperiod-sensitive and -insensitive maize lines

*GI* is specific to terrestrial plants whose expression is regulated by the circadian clock, and it functions as a regulator of biological rhythms and flowering in plants (Imaizumi et al 2005, Park et al 2013). The circadian clock system is regulated in an orderly and precise manner by multiple interconnected transcriptional and translational feedback loops. The *GI* repressors *CCA1* and *LHY* are inhibited during the day by *TOC1*, which is in turn degraded at night by the F-box protein *ZEITLUPE* (ZTL), releasing *CCA1* and *LHY* to inhibit *GI* expression (Cha et al 2017, David et al 2006, Más et al 2003), and thus resulting in the *GI* rhythmic expression pattern. The FPKM and qPCR analyses showed that *ZmGI1* expression follows a typical circadian rhythmic pattern, reaching peak expression in the evening near the onset of darkness and a lowered expression at dawn. This indicates that the regulation of *ZmGI1* expression depends highly on photoperiod. *ZmGI1* mRNA accumulates in a similar fashion to that of *TOC1* and *CO*, which enhance flowering under prolonged photoperiodic conditions (Doyle et al 2002, Más et al 2003). We believe that the circadian rhythmic expression of *ZmGI1* probably resulted from the accumulation and inhibition of the ZmGI1 protein during the day and night, respectively.

*GI* genes are known to function in the photoperiodic pathway as part of a timekeeping mechanism that regulates the perception of photoperiodic cues by photoperiod-sensitive plants (Mizoguchi et al 2005, Samach & Coupland 2000). The results of the expression analysis showed marked variation in the rhythmic expression of *ZmGI1* under LD conditions in the different maize inbred lines. The tropical sensitive lines exhibited higher *ZmGI1* expression and reached peak expression 3 h earlier than the temperate insensitive inbred lines. Moreover, the expression of *ZmGI1* among the lines was significantly and positively correlated with DTS under LD conditions (r = 0.91, *P* < 0.05) (Figure 2C); this implies that *ZmGI1* plays a central role in regulating the sensitivity of tropical maize lines to LD photoperiodism compared with the insensitive lines, and thus its regulatory role differs between the tested tropical and temperate lines.

### ZmGI1 plays a crucial role in regulating flowering time

The results of this study revealed that *ZmGI1* exhibits typical circadian characteristics and regulates flowering under LD conditions. SNP variations of *ZmGI1* exhibited significant associations with flowering-related traits in maize, while overexpression in Arabidopsis significantly promoted flowering under LD conditions compared to the WT and *gi* mutants. The complex of ZmGI1 and ZmFKF1a can regulate the degradation of CO and FT repressors under LD conditions in Arabidopsis. Mizoguchi (Mizoguchi et al 2005) reported late flowering and a reduction in *CO* mRNA accumulation in *gi* mutants, and suggested that *GI* might function in the flowering pathway in Arabidopsis by regulating a surge in the abundance of mRNAs of *CO* and *FT* under LD conditions. Accumulation of CO and promotion of flowering in overexpressing *ZmGI1* (Figure 3) was in tandem with the findings of Imaizumi *et al*. (Imaizumi et al 2005) and Suarez-Lopez (Suárez-López et al 2001). Since CO and FT proteins are highly conserved among photoperiod-sensitive plants (Ballerini & Kramer 2011), *ZmGI1* is believed to act upstream and activates the transcription of *CO* and *FT*, which function as central integrators for regulating flowering time in photoperiod-sensitive plants. At the posttranscriptional stage, the stability of CO at the end of the light period under LD conditions is modulated by the circadian clock through the regulation of *GI* and *FKF1* expression (Park et al 2016, Song et al 2015). In addition, comparisons of the bolting time and rosette leaves among the genotypes showed a significant delay in flowering time and higher number of rosette leaves in the *gi* mutants compared to the overexpression lines and WT, confirming the role of *ZmGI1* in the flowering pathway in plants.

However, the flowering regulatory function of *ZmGI1* was widely divergent in Arabidopsis and in diverse maize germplasms. The role of *ZmGI1* in regulating circadian rhythms, revealed in the expression analysis, might have contributed to its function in the photoperiodic regulation of maize flowering time. This is consistent with the findings made by Li *et al*. (Li et al 2016) and Salomé *et al*. (Salomé et al 2008). The expression and correlation analyses also demonstrated that *ZmGI1* is functionally distinct in various maize inbred lines under LD and SD photoperiodic conditions. The abundance of *ZmGI1* transcripts was positively correlated with days to flowering time in the three tropical lines under LDs, and the *ZmGI1* expression level was positively correlated with DTS, suggesting an inhibiting effect of *ZmGI1* on regulating flowering time under LD conditions.

### Dual functions of ZmGI1 in flowering time regulation and salt stress response

*ZmGI1* expression was revealed in roots, stems, leaves, floral organs, and fruit spikes, and we found that it could regulate other biological functions besides the circadian clock and plant flowering pathways. *GI* has been implicated in pleotropic functions, including perception and adaptation to environmental stress conditions (Kim et al 2013, Mishra & Panigrahi 2015, Riboni et al 2013). Differences in the function of different mutants generated from mutations at different segments of *GI* indicate that different segments of *GI* might have distinct or opposing regulatory roles in plants. A similar deduction was made by Kim *et al*. (Kim et al 2016), who used RNAi technology to selectively degrade different segments of *BrGI* and generated different mutants with opposing responses to salt stress.

In this study, we have demonstrated that *ZmGI1* has multiple functions in maize and that there were linkages between the regulation of the photoperiodic flowering pathway and salt stress response (Figure 4). In the salt response, an anticipated accumulation of proline, which helps to enhance stress tolerance in plants, was detected in both roots and leaves (Figure S4A). H_2_O_2_, produced by cellular aerobic metabolism, seems to be tightly regulated as H_2_O_2_ levels increase only slightly in response to stress (Rani et al 2015). By contrast, H_2_O_2_ has been found to act as a signaling molecule and secondary messenger to regulate various stress resistance processes in plant (Rani et al 2015, Sewelam et al 2014). Thus, the balance of H_2_O_2_ in cells maintained by the antioxidant enzymes was an important indicator in salt stress response. In this study, NaCl promoted the accumulation of H_2_O_2_ in roots. Low levels of H_2_O_2_ act as a signal to activate the expression of ZmThox, a member of the thioredoxin peroxidases, which then eliminates the H_2_O_2_ in leaves to reduce damage to leaves from oxygen free radicals (Figure 8). In addition, ZmThox also belongs to the PPPDE family, a kind of deubiquitinating enzyme, and inhibits the degradation of ZmGI1 under salt stress.

**Figure 8.**
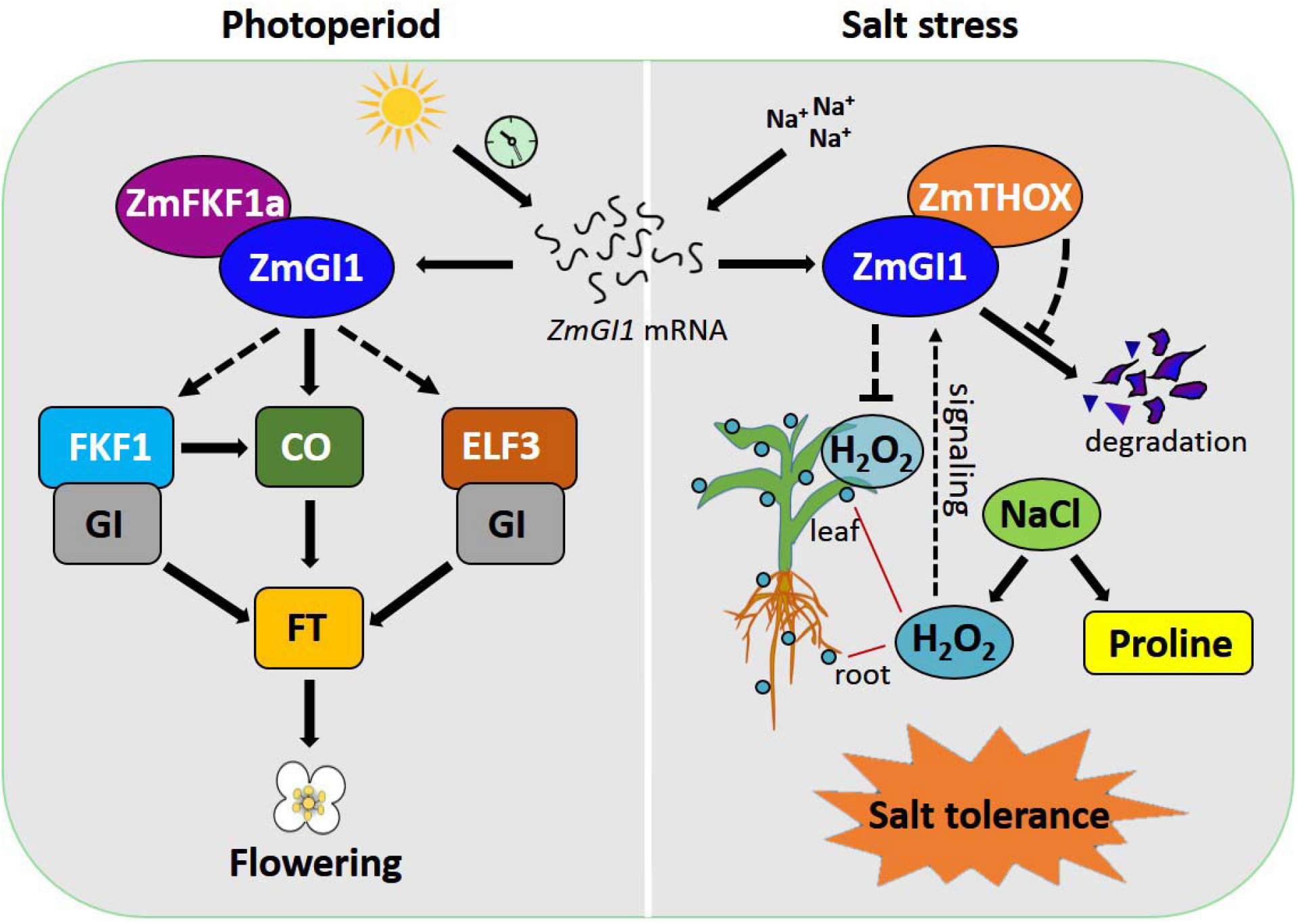
A proposed working model of *ZmGI1-*mediated flowering promoter and salt resistance. The photoperiodic flowering pathway is on the left. ZmGI1 interacts with ZmFKF1a and promotes flowering through enhancing *CO* expression. Overexpression of *ZmGI1* up-regulates the expression of downstream genes including *FKF1, ELF3* and *CO*, which will result in accumulation of florigen and promoted flowering in *Arabidopsis*. ZmGI1 interacts with ZmTHOX, a component of redox balance pathway is on the right, which belongs to PPPDE peptidase family involved in substrate deubiquitinating. Salt stress (with NaCl) triggers the up-regulation of ZmThox under the stimulation of H_2_O_2_ in roots, then act on the stabilization of ZmGI1and the elimination of H_2_O_2_ in leaves. While at the transcriptional level, photoperiodic-insensitive lines have low *ZmGI1* expression. Salt stress resulting in the accumulation of proline in plant that enhance salt stress tolerance. The black solid arrow indicates that the data is from this study.

We concluded that overexpression of *ZmGI1* confers salt stress tolerance in Arabidopsis, which is contrary to the findings of Kazan and Lyons (Kazan & Lyons 2016), who reported that *AtGI* negatively regulates salt tolerance by repressing the phosphorylation of SOS1 in the SOS pathway. The ZmGI1/ZmThox complex might confer salt stress tolerance by protecting the mitochondria from oxidative stress through a post-transcriptional adjustment in *S*-glutathionylation and *S*-nitrosylation. The role of the *Trx* gene family in oxidative stress tolerance was elaborated by Lázaro *et al*. (Lázaro et al 2013). Generally, we found that ZmGI1 had significant roles in the photoperiodic flowering pathway and salt stress in maize. However, the molecular mechanisms underlying the function of ZmGI1 are likely to be widely divergent from GI functions in Arabidopsis. Studies exploring whether ZmGI1 has even more functional roles should be conducted, and the molecular mechanisms of ZmGI1 function in different cellular processes should be further investigated.

## Supplementary data

**Figure S1** The expression characteristics of *ZmGI1*.

**Figure S2** Expression pattern of flowering relative genes in WT, gi and overexpression plants under LDs.

**Figure S3** Pro35S::ZmGI1 affect plant growth responses to salt stress in Arabidopsis thaliana.

**Figure S4** Determination of proline and H_2_O_2_ content under salt stress in maize seedlings.

**Figure S5** Transcriptional activity assay of ZmGI1.

**Table S1** Primers used in this study.

**Table S2** Associations between the natural variations within ZmGI1 and agronomic traits.

## Acknowledgment

We are grateful to Prof. Cuijun Zhang (Shenzhen Agricultural Genome Research Institute, Chinese Academy of Agricultural Sciences) and Prof. Chunzhao Zhao (Institute of Plant Physiology and Ecology, Chinese Academy of Sciences) for technical assistance. This work was supported by grants from the National Key Research and Development Program of China (2016YFD0101803); and the National Natural Science Foundation of China (32030078 and 31901557).

## Author Contributions

Y.L., F.W. and L.L. conceived of the research, and participated in its design and coordination; F.W., L.L., and Y.K. performed the experiments and drafted the manuscript; J.L., Z.M., Y.K., B.S.Y., and E.H. performed data collection, analysis, or interpretation; J.X., Q.W., X.F. and Y.L. revised the manuscript. F.W., L.L., and J.L. contributed equally. All the authors read and approved of the final version of the manuscript.

## Notes

### Competing Interest Statement

The authors have declared no competing interest.

